# QTL analysis of macrophages from an AKR/JxDBA/2J intercross identified the *Gpnmb* gene as a modifier of lysosome function

**DOI:** 10.1101/684118

**Authors:** Peggy Robinet, Brian Ritchey, Alexander M. Alzayed, Sophia DeGeorgia, Eve Schodowski, Shuhui Wang Lorkowski, Jonathan D. Smith

## Abstract

Our prior studies found differences in the AKR/J and DBA/2J strains in regard to atherosclerosis and macrophage phenotypes including cholesterol ester loading, cholesterol efflux, and autolysosome formation. The goal of this study was to determine if there were differences in macrophage lysosome function, and if so to use quantitative trait locus (QTL) analysis to identify the causal gene. Lysosome function was measured by incubation with an exogenous double-labeled ovalbumin indicator sensitive to proteolysis. DBA/2J vs. AKR/J bone marrow macrophages had significantly decreased lysosome function. Macrophages were cultured from 120 mice derived from an AKR/JxDBA/2J F_4_ intercross. We measured lysosome function and performed a high density genome scan. QTL analysis yielded two genome wide significant loci on chromosomes 6 and 17, called macrophage lysosome function modifier (*Mlfm*) loci *Mlfm1* and *Mlfm2*. After adjusting for *Mlfm1*, two additional loci were identified. Based on proximity to the *Mlfm1* peak, macrophage mRNA expression differences with AKR/J >> DBA/2J, and a protein coding nonsense variant in DBA/2J, the *Gpnmb* gene, encoding a lysosomal membrane protein, was our top candidate. To test this candidate, *Gpnmb* expression was knocked down with siRNA in AKR/J macrophages; and, to express the wildtype *Gpnmb* in DBA/2J macrophages, we obtained a DBA/2 substrain, DBA/2J-Gpnmb+/SjJ, which was isolated from the parental strain prior to its acquiring the nonsense mutation, and subsequently back crossed to the modern DBA/2J background. Knockdown of *Gpnmb* in AKR/J macrophages decreased lysosome function, while restoration of the wildtype *Gpnmb* allele in DBA/2J macrophages increased lysosome function. However, this modifier of lysosome function was not responsible for the strain differences in macrophage cholesterol ester loading or cholesterol efflux. In conclusion, we identified the *Gpnmb* gene as the major modifier of lysosome function and we showed that the ‘QTL in a dish’ strategy is efficient in identifying modifier genes.

**Author Summary:** Inbred strains of mice differ in both their genetic backgrounds as well as in many traits; and, classical mouse genetics allows the mapping of genes responsible for these traits. We identified many traits that differ between the inbred strains AKR/J and DBA/2J, including atherosclerosis susceptibility, macrophage cholesterol metabolism, and in the current study, macrophage protein degradation via an organelle called the lysosome. Using mouse genetic mapping and bioinformatics we identified a candidate gene, called *Gpnmb*, responsible for modifying lysosome function; and, the DBA/2J strain carries a mutation in this gene. Here we demonstrate that the *Gpnmb* gene is a modifier of lysosome function by either correcting this *Gpnmb* mutation in DBA/2J macrophages, or by knocking down *Gpnmb* expression in AKR/J macrophages. This study is noteworthy as the human *GPNMB* gene has been implicated in many diseases including cancer, kidney injury, obesity, non-alcoholic steatohepatitis, Parkinson disease, osteoarthritis, and lysosome storage disorders.

## Introduction

Atherosclerosis, the primary cause of cardiovascular disease and the leading cause of death worldwide[1], is characterized by the progressive buildup of cholesterol-rich plaques in the arteries. Atherosclerosis severity in various mouse models is modified by their genetic background[2]. On the apoE-deficient background, AKR/J mice have 10-fold smaller aortic root lesions than DBA/2J[2]. When bone marrow derived macrophages (BMDM) from these two strains are loaded with acetylated low density lipoprotein (AcLDL) in vitro, modeling cholesterol loaded foam cells, DBA/2J cells accumulate more cholesterol esters (CE) while AKR/J cells have higher free cholesterol (FC) levels leading to CE/FC ratio ~3-fold higher in DBA/2J macrophages[3]. Autophagy is the major pathway for lipid droplet clearance in these foam cells. This involves the engulfment of lipid droplets into autophagosomes, which fuse with lysosomes where CE is hydrolyzed to FC by lysosomal acid lipase. DBA/2J foam cells have delayed autolysosome formation resulting in an inefficient clearance of lipid droplet CE[3]; whereas, autophagy initiation and autophagosome number is not different in AKR/J vs. DBA/2J foam cells[3]. We thus made the hypothesis that the lysosome arm of autophagy, rather than the autophagosome arm, was leading to the observed differences in cholesterol metabolism in AKR/J vs. DBA/2J macrophage foam cells.

Here, we applied a genetic instrument, quantitative trait locus (QTL) mapping, to identify genes that impact lysosome function in BMDM from an AKR/JxDBA/2J F_4_ strain intercross. We discovered four macrophage lysosome function modifier (*Mlfm*) loci, with the strongest locus on the proximal region of chromosome 6 (*Mlfm1*). The gene encoding the Glycoprotein Non-Metastatic Protein B (*Gpnmb*) was our top candidate gene for the *Mlfm1* QTL due to its proximity to the LOD peak, expression difference and expression QTL (eQTL) in our strain pair[4, 5], and the presence of a nonsense mutation in the DBA/2J strain[6]. In the context of macrophages, the GPNMB protein has been shown to be associated with lysosome and autophagosome membranes[7–9]. *Gpnmb* expression is increased in foam cells[4, 10], especially M2 polarized macrophages[11]. This GPNMB protein is also induced in lysosomal storage diseases[12]. We now describe our findings that the *Gpnmb* gene is responsible for the *Mlfm1* QTL. However, the effect of the *Gpnmb* gene on lysosome function did not translate to an effect on cholesterol ester loading or cholesterol efflux. Thus, we still have not discovered the gene responsible for decreased lysosome-autophagosome fusion in DBA/2J BMDM that can alter lipid droplet turnover.

## Results

### DBA/2J vs. AKR/J BMDM have decreased lysosome function

We assessed lysosome volume per cell by immunofluorescent staining of fixed cells with Lamp1 antibody and flow cytometry. DBA/2J vs. AKR/J BMDM had a 32% decrease in lysosome volume (p<0.01, Figure 1A). In order to measure lysosome function in AKR/J and DBA/2J BMDM, we adopted a commercially available compound, DQ-ovalbumin. This product is taken up by cellular pinocytosis and accumulates in lysosomes, where proteolysis increases the fluorescence of the tightly-packed self-quenched Bodipy fluorophore[13]. We showed that cellular Bodipy fluorescence was blunted by pretreatment of the cells with lysosomal protease inhibitors E64d and pepstatin A, which also get into cells via pinocytosis (Figure 1B). As fluorescence intensity would also be dependent upon the uptake of the label, we modified the DQ-ovalbumin by covalent modification with Alexa647 (A-DQ-ova), so that the Bodipy/Alexa647 fluorescence ratio would indicate proteolysis normalized for cellular uptake. We validated this doubly labeled A-DQ-ova by incubation with proteinase K, which showed robust increase in the Bodipy/Alexa647 fluorescence ratio (Figure 1C). We assessed lysosome function by flow cytometry and calculated the Bodipy/Alexa647 fluorescence intensity ratio in 10,000 cells per line (Figure 1D). The median Bodipy/Alexa647 ratio was 45% higher in in AKR/J vs. DBA/2J (p<0.01, Figure 1E left panel), while the 95^th^ percentile was 49% higher in AKR/J vs. DBA/2J (p<0.001, Figure 1E right panel); demonstrating decreased lysosome function in DBA/2J BMDM.

**Figure 1.**
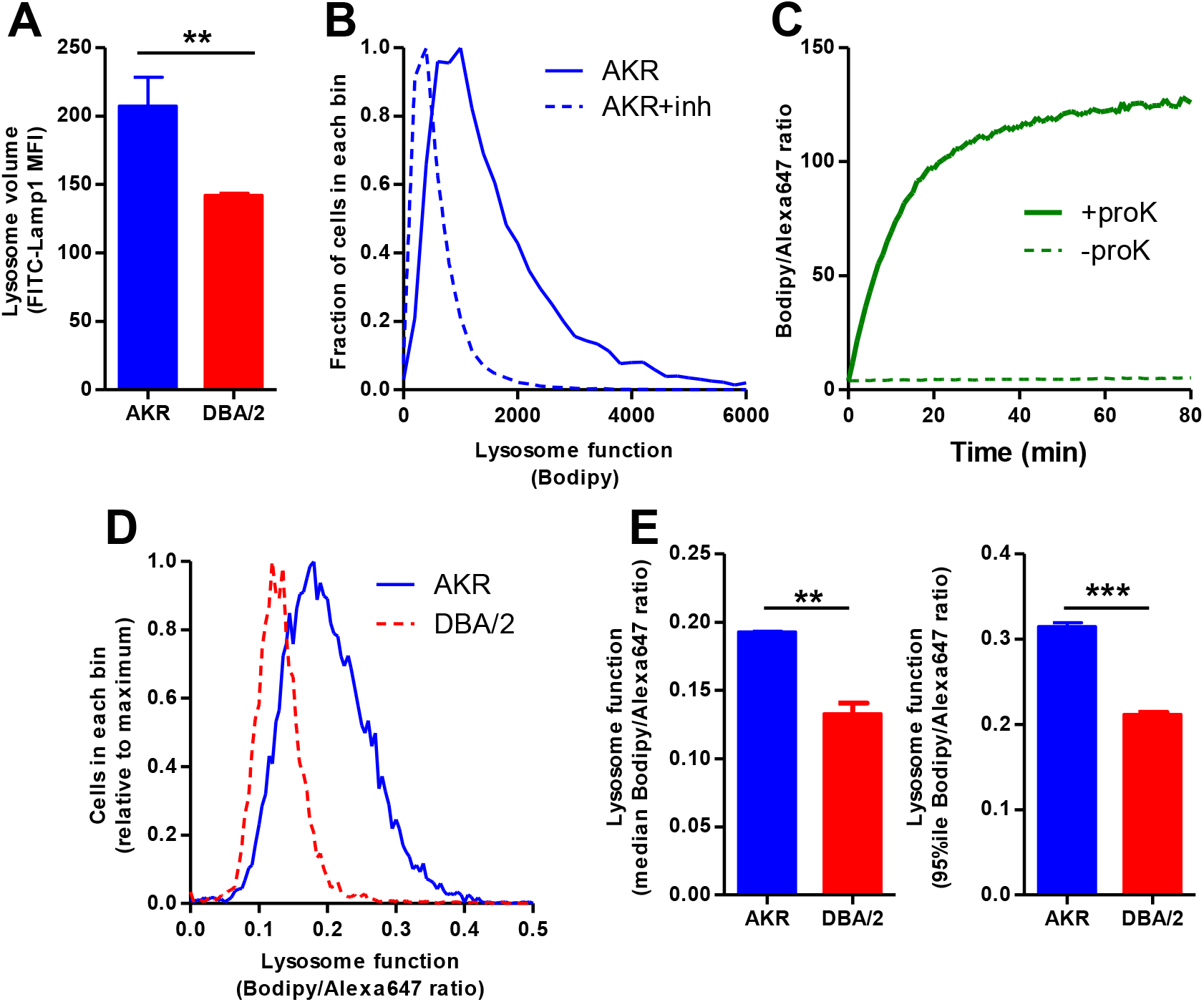
Decreased lysosome function in DBA/2J vs. AKR/J macrophages. **A.** Lysosome volume assessed by Lamp1 immunofluorescence and flow cytometry (median fluorescence intensity; n=3 per strain, AKR blue bars; DBA/2 red bars). **B.** Lysosome function in AKR BMDM assessed by incubation with DQ-ovalbumin and flow cytometry (solid line). Lysosome inhibition with E64d plus pepstatin A shows a leftward shift (dashed line) indicating decreased lysosome function. **C.** In vitro validation of A-DQ-ova reagent showing proteinase K increases Bodipy/Alexa647 fluorescence ratio. **D.** Typical lysosome function assay in AKR (solid blue line) and DBA/2 (dashed red line) BMDM assessed by incubation with A-DQ-ova and flow cytometry. **E.** Analysis of lysosome function by A-DQ-ova incubation in AKR (blue bars) and DBA/2 (red bars) using median (left panel) or the 95^th^ percentile (right panel) fluorescence intensity ratio (duplicate assay). **, p<0.01; ***, p<0.001 by two-tailed t-test.

We then incubated cells with FITC-TAMRA-dextran to assess lysosomal pH by flow cytometry. Dextran is taken up by pinocytosis and accumulates in lysosomes. FITC fluorescence is pH dependent and is lower as pH decreases in acidic organelles such as lysosomes, while TAMRA fluorescence is pH insensitive. Thus, cellular FITC/TAMRA ratio is an indicator of lysosomal pH. We demonstrated that treatment of cells with Bafilomycin A1 led to a time dependent increase in the FITC/TAMRA ratio, indicating increased lysosomal pH (Figure 2A). We calculated FITC/TAMRA ratio in AKR/J vs. DBA/2J BMDM and found that the lysosomal pH was not statistically different (Figure 2B, C), ruling out increased pH as the cause for lysosome dysfunction in DBA/2J macrophages.

**Figure 2.**
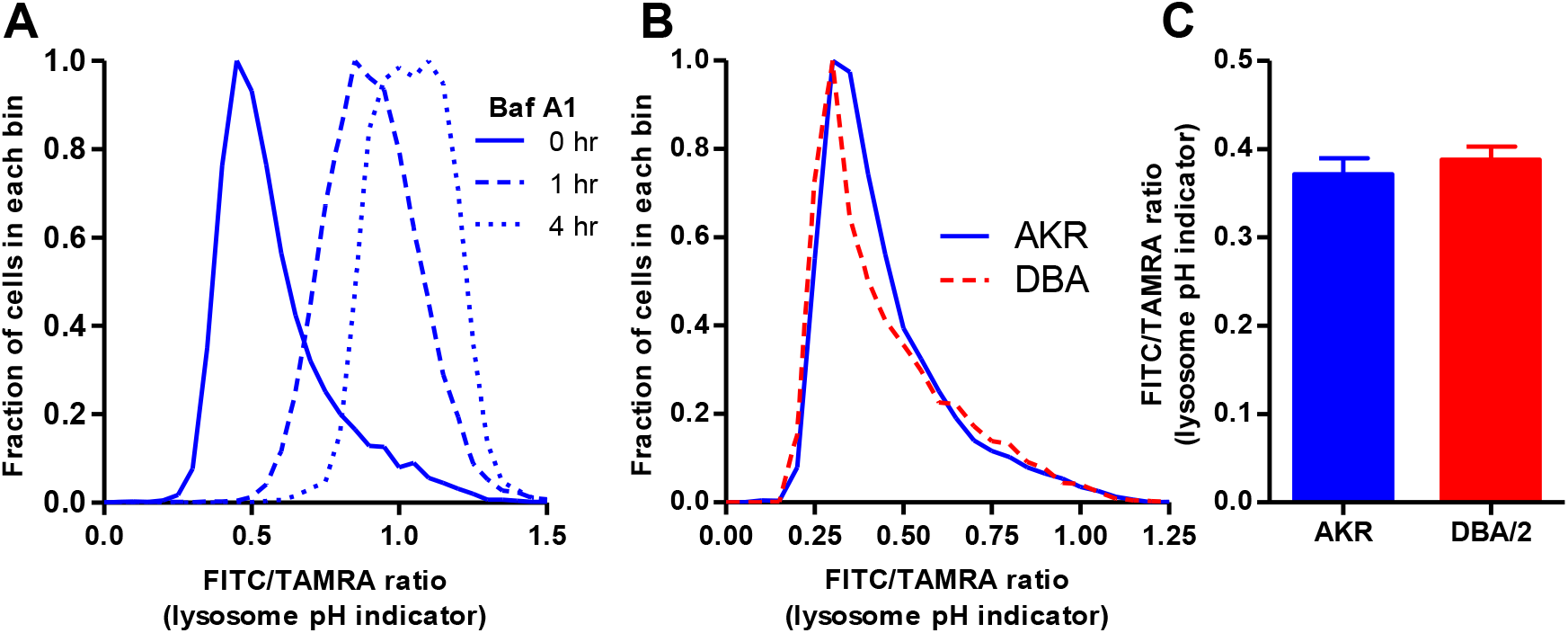
Unaltered lysosomal pH in DBA/2J vs. AKR/J macrophages. **A.** AKR macrophages were incubated for 18h with TAMRA-FITC dextran. This was followed by a 4h equilibration -/+ Bafilomycin A1 added for the indicated times before cell harvest. Cells were analyzed by flow cytometry, demonstrating the usefulness of this probe to assess lysosomal pH. **B.** Typical lysosomal pH assay in AKR (solid blue line) and DBA/2 (dashed red line) BMDM assessed by incubation with TAMRA-FITC dextran and flow cytometry. **C.** Analysis of lysosomal pH in AKR (blue bars) and DBA/2 (red bars) using median fluorescence intensity ratio (n=3, not significant by two tailed t-test).

### Significant QTL for lysosome function maps to chromosome 6

We used a genetic approach to identify the gene region responsible for the strain difference in lysosome function. Lysosome function was measured using A-DQ-ova in BMDM lines derived from 120 F_4_ mice from an AKR/JxDBA/2J strain intercross, using the 95^th^ percentile values, as these were more significant than the median values (Figure 1E). There was no significant effect of sex on this phenotype (p=0.61), so we combined data from male and female macrophages. The data were normally distributed (Figure 3A) and were used to perform QTL mapping. We identified two macrophage lysosome function modifier (*Mlfm*) QTLs that we named *Mlfm1* and *Mlfm2* on the proximal regions of chromosomes 6 and 17, respectively (Figure 3B). *Mlfm1*, at 49.7 Mb on chromosome 6 (90% confidence interval 28.7 – 64.9 Mb), was the strongest locus with a LOD score of 6.09 (Table 1). We divided the 120 BMDM lines by their genotypes at *Mlfm1* and found a gene dosage effect with each DBA/2J allele decreasing lysosome function by 6% (Figure 3C, ANOVA linear trend test r^2^=0.208, p<0.0001), indicating that this locus is associated with ~21% of the variance in lysosome function in the F_4_ cohort. *Mlfm2* was located at 9.5 Mb on chromosome 17 (90% confidence interval 6.0 – 16.5 Mb) with a LOD score of 4.28 (Table 1). We adjusted for the *Mlfm1* genotype as an additive co-variate and reran the QTL analysis (Figure 4). We identified new peaks on the distal sides of chromosomes 2 and 17 (*Mlfm3* LOD score 5.55 and *Mlfm4* LOD score 4.32, respectively) and confirmed *Mlfm2* on the proximal end of chromosome 17 that was moderately strengthened (LOD score 4.61, Table 2).

**Figure 3.**
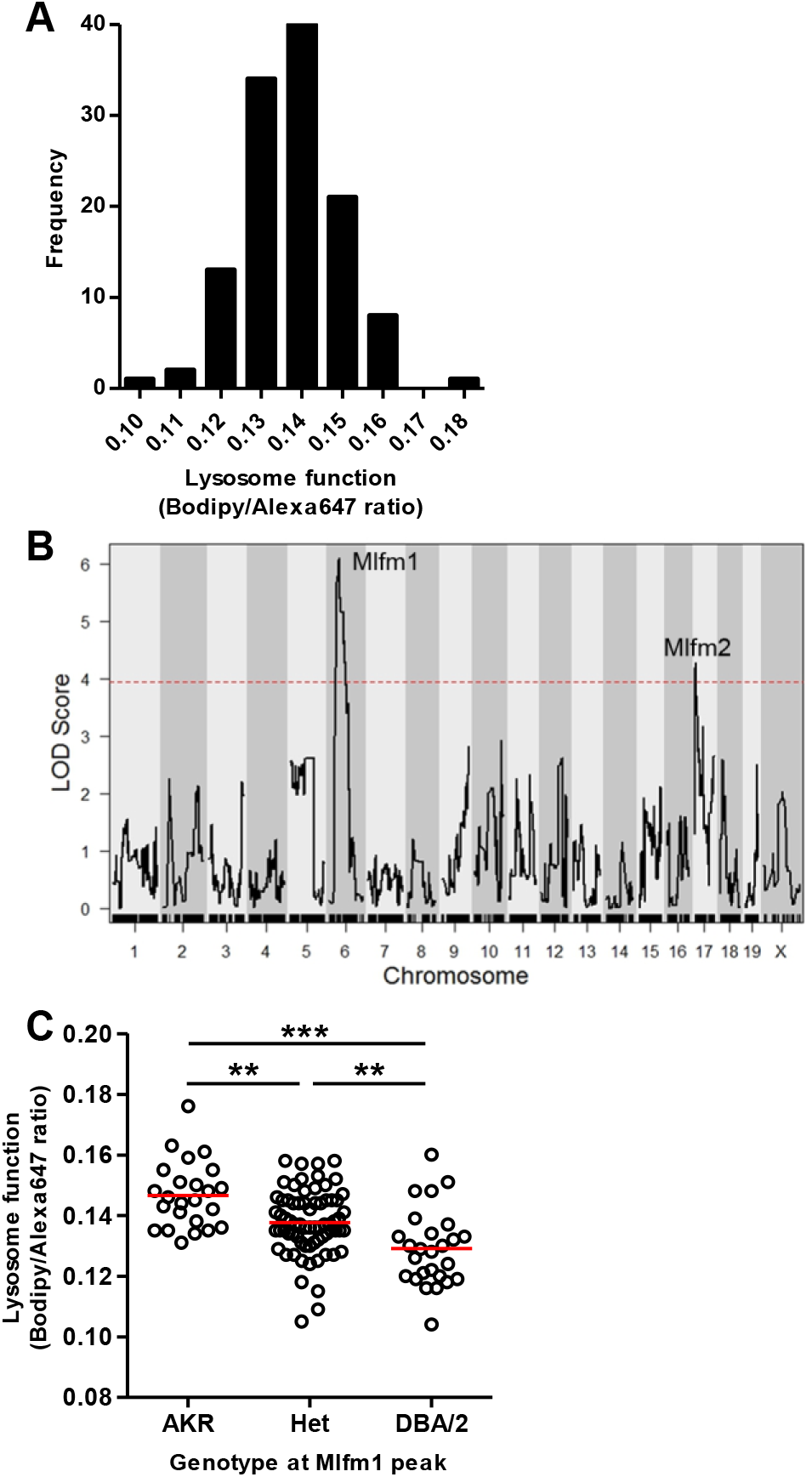
QTL analysis for lysosome function. **A.** Normal distribution of the Bodipy/Alexa647 fluorescence ratio after F_4_ BMDM incubation with A-DQ-ova. **B.** QTL LOD plot for lysosome function showing *Mlfm1* and *Mlfm2* on chromosomes 6 and 17, respectively. The dashed red line shows the genome-wide p=0.05 threshold based on 10,000 permutations. **C.** F_4_ BMDM lysosome function values by genotype at the *Mflm1* peak marker (mean values (red lines); ANOVA linear trend test r^2^=0.208, p<0.0001; **, p<0.01, and ***, p<0.001 by ANOVA Tukey posttest).

**Table 1.**
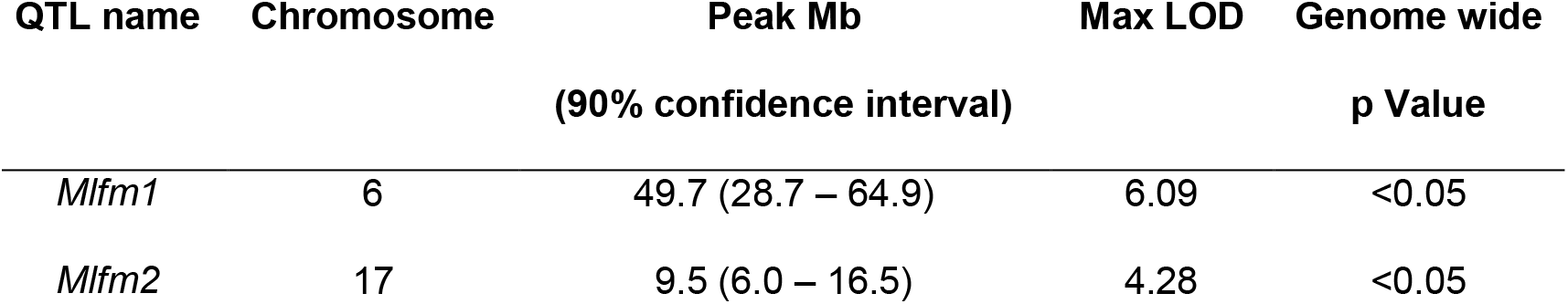
Significant lysosome function QTLs

**Figure 4.**
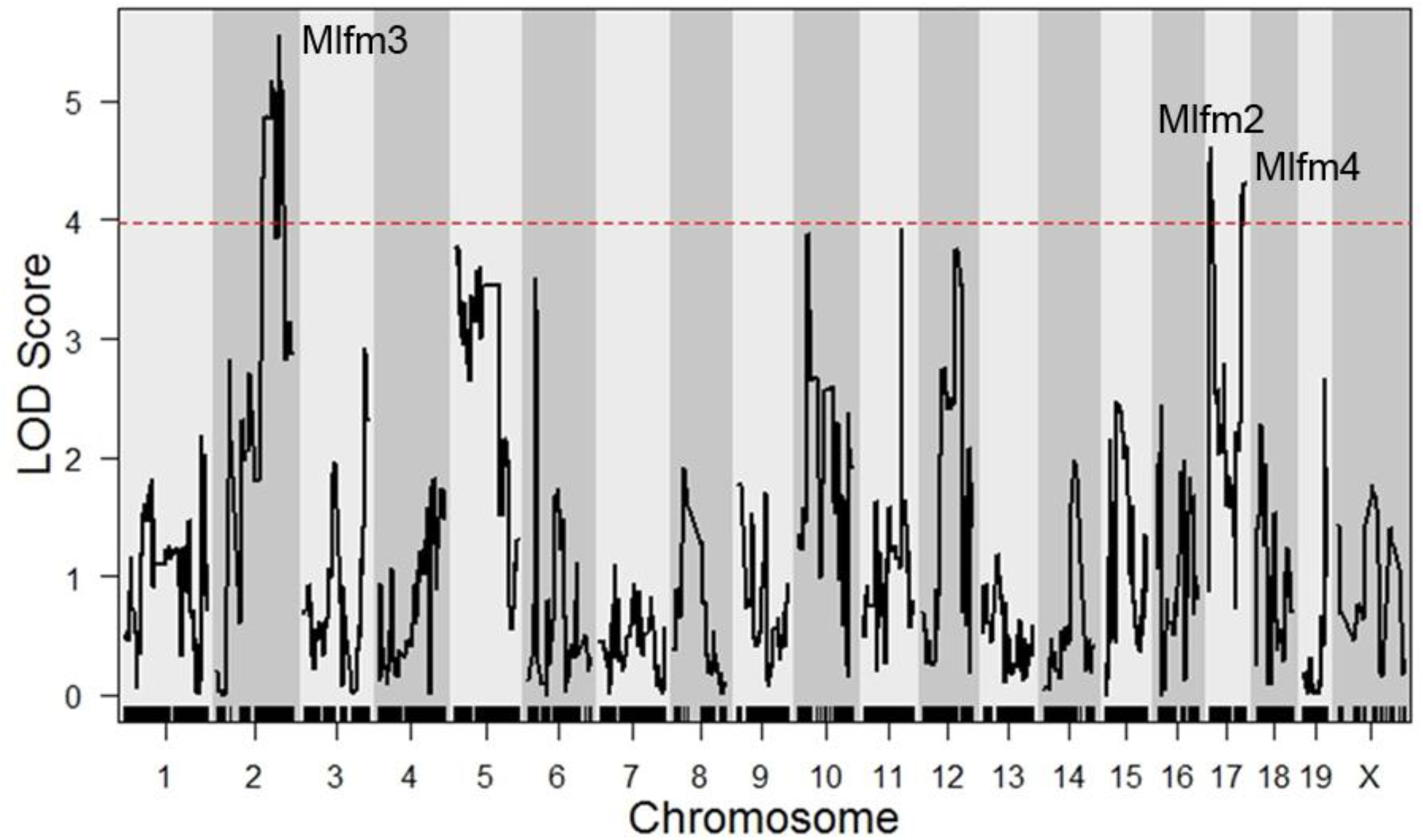
QTL analysis for lysosome function after adjusting for *Mlfm1* as an additive co-variate. The dashed red line shows the genome-wide p=0.05 threshold based on 10,000 permutations.

**Table 2.** Significant lysosome function QTLs after adjusting for *Mlfm1*.

| QTL name | Chromosome | Peak Mb<br>(90% confidence interval) | Max LOD | Genome wide<br>p Value |
| --- | --- | --- | --- | --- |
| *Mlfr2* | 17 | 9.6 (6.0 – 17.1) | 4.61 | <0.05 |
| *Mlfr3* | 2 | 147.5 (111.2 – 158.1) | 5.55 | <0.05 |
| *Mlfr4* | 17 | 87.5 (79.4 – 89.1) | 4.32 | <0.05 |

### Identification of *Gpnmb* as the gene responsible for the *Mlfm1* QTL

To limit the *Mlfm1* locus further we performed a Bayesian analysis which yielded an interval of 25.89 to 63.78 Mb, which contained 430 genes. Among them, *Gpnmb*, mapping at 48.99 Mb (0.17 Mb from the LOD peak), has a C>T mutation in DBA/2J mice leading to an early stop codon in exon 4[6]. This mutation is predicted to lead to non-sense mediated mRNA decay as traditionally defined[14]. We had previously observed a DBA/2J-AKR/J strain difference in BMDM *Gpnmb* mRNA levels in a microarray study with DBA/2J macrophages expressing ~12.5 fold less *Gpnmb* mRNA[4]. In a prior independent F_2_ strain intercross we found a strong cis expression QTL (eQTL) for *Gpnmb* expression in BMDM with a LOD score of 22[5]. We performed a western blot for GPNMB protein in lysates from AKR/J and DBA/2J BMDM, and we did not detect any expression in DBA/2J macrophages (Figure 5A). Thus, we prioritized the *Gpnmb* gene as our top candidate at this locus. Fortunately, there exists a DBA/2 substrain available at JAX that contains the wildtype *Gpnmb* gene, called DBA/2J-Gpnmb^+^/SjJ, which we will refer to as DBA/2g^+^[15]. Western blot confirmed GPNMB protein expression in this line (Figure 5A). We used siRNA to knockdown *Gpnmb* in AKR/J BMDM, referred to as AKRg^-^, which decreased GPNMB protein expression robustly (Figure 5A).

**Figure 5.**
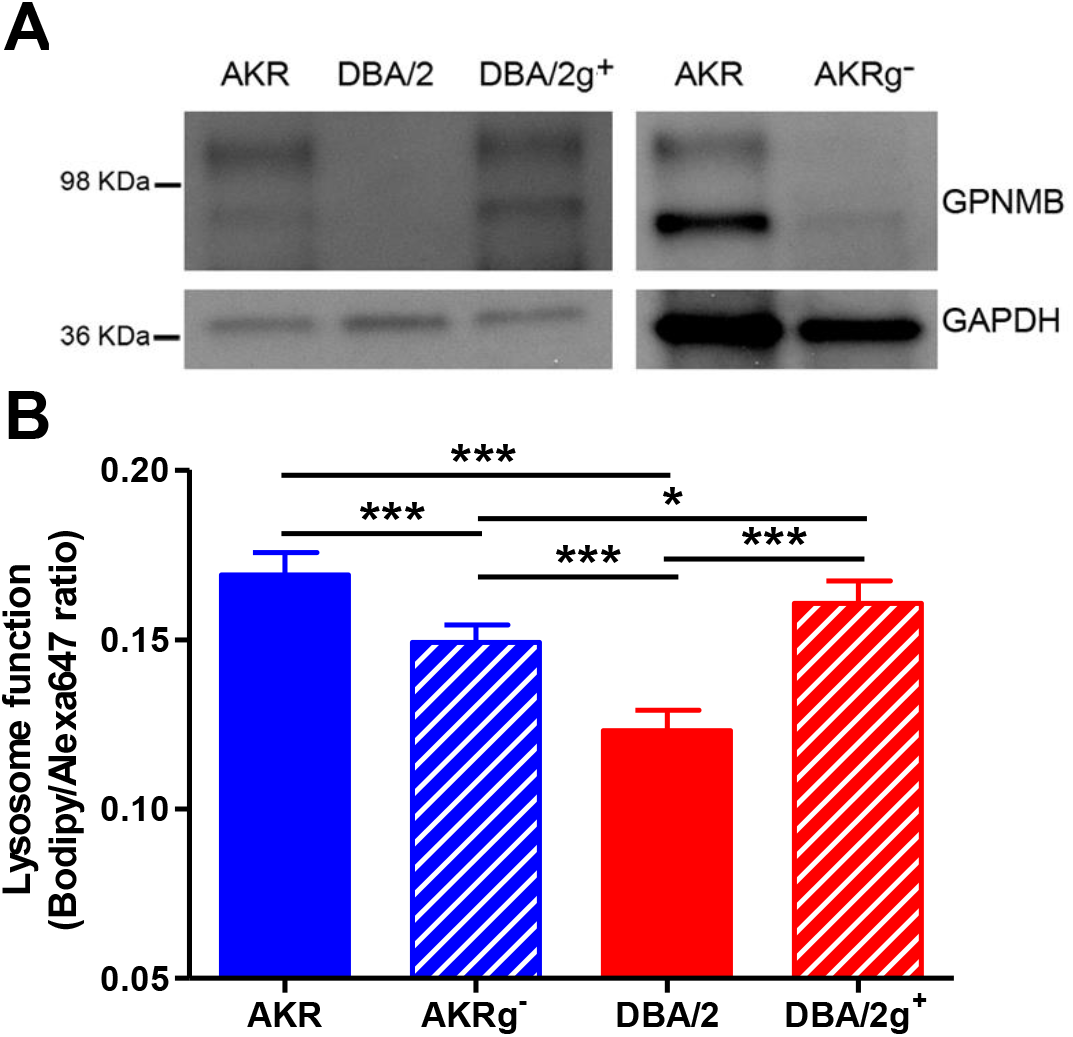
Altered lysosome function dependent upon *Gpnmb* expression. **A.** GPNMB and GAPDH western blot from lysates of AKR, DBA/2, and DBA/2g+ BMDM (left panel); and from AKR BMDM transfected with control or *Gpnmb* siRNA (right panel). **B.** Analysis of lysosome function by A-DQ-ova incubation in BMDM from AKR control siRNA (solid blue bars), AKRg^-^ (striped blue bars), DBA/2 (solid red bars), and DBA/2g^+^ (striped red bars) using median fluorescence intensity ratio (n=5; *, p<0.05; ***, p<0.001 by ANOVA with Tukey posttest).

Lysosome function was measured in AKR/J, AKRg^-^, DBA/2J and DBA/2g^+^ BMDM using A-DQ-ova (Figure 5B). DBA/2J vs. AKR/J BMDM had a 27% decrease in lysosome function (p<0.001, by ANNOVA posttest). AKRg^-^ vs. AKR/J had a 12% decrease in lysosome function (p<0.001). WT *Gpnmb* expression in DBA/2g^+^ restored lysosome function to level similar than those observed in AKR/J BMDM (DBA2/g^+^ vs. DBA/2J, 30% increase, p<0.001; DBA/2g^+^ vs. AKR/J, not significant). These data confirm *Gpnmb* as a causal gene at the *Mlfm1* locus, the strongest locus associated with lysosome function.

### *Gpnmb* expression does not alter macrophage cholesterol loading or efflux

We tested to see if *Gpnmb* genotype altered cholesterol loading and efflux. The 4 genotypes of BMDM were loaded with 50 μg/mL AcLDL for 24h. Total cholesterol levels were not significantly different, but as we had previously observed[3, 16], AKR/J cells accumulated more free cholesterol and DBA/2J cells accumulated more cholesterol esters (Figure 6 A-C). However, *Gpnmb* gene status had no effect on cholesterol loading, only strain effects were significant. [^3^H]Cholesterol labeled loaded BMDM were allowed to efflux cholesterol for 4h to 10 μg/mL lipid-free apoA1. Again, *Gpnmb* gene status did not alter efflux, only the strain effect was significant as previously described[3].

**Figure 6.**
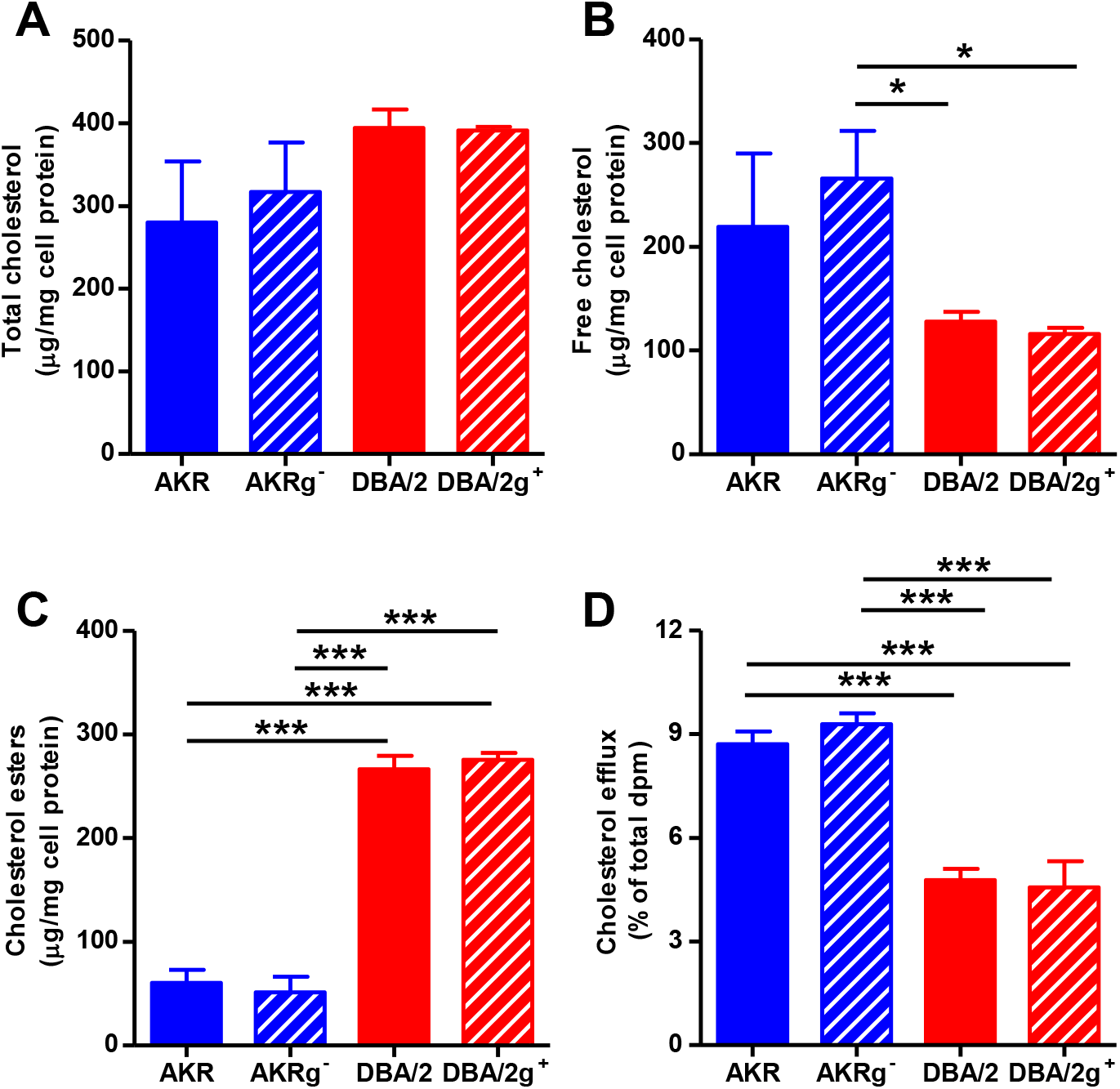
Cholesterol loading and efflux independent of *Gpnmb* expression. **A-C.** Total cholesterol level (A), free cholesterol (B) and cholesterol esters (C) normalized to protein levels in AcLDL loaded BMDM from AKR control siRNA (solid blue bars), AKRg^-^ (striped blue bars), DBA/2 (solid red bars), and DBA/2g^+^ (striped red bars) (n=3; *, p<0.05; ***, p<0.001 by ANOVA with Tukey posttest). **D.** Efflux of cholesterol to lipid-free apolipoprotein A1 from AcLDL loaded BMDM from AKR control siRNA (solid blue bars), AKRg^-^ (striped blue bars), DBA/2 (solid red bars), and DBA/2g^+^ (striped red bars) (n= 3; ***, p<0.001 by ANOVA with Tukey posttest).

## Discussion

We previously crossed apoE-deficiency onto six inbred strains, DBA/2J, C57BL/6J, 129/SV-ter, AKR/J, BALB/cByJ, and C3H/HeJ; and, among these strains the DBA/2J has the largest aortic root atherosclerotic lesions, while AKR/J was one of several strains with small lesions[2]. This led us to follow up with two independent strain intercrosses to identify atherosclerosis modifier genes using the DBA/2J and AKR/J parental strains, which identified three confirmed atherosclerosis QTLs, *Ath22*, *Ath26*, and *Ath28*[5, 17]. Since macrophages are a key cell type in atherogenesis, we also performed eQTL analysis of BMDM from these same two independent AKR/JxDBA/2J strain intercrosses in order to gain insights into potential atherosclerosis modifier candidate genes[5, 18]. We also started series of studies to explore BMDM phenotypes from these two parental strains. Thus far we found significant strain effects on cholesterol ester loading, cholesterol efflux to apolipoprotein AI or HDL acceptors, autolysosome formation, and, in the present study, lysosome function. To identify QTL loci for these traits, we bred an F_4_ strain intercross and froze down aliquots of bone marrow for subsequent phenotype studies. We recently reported the first of these QTL studies, which identified an AKR/J deletion in exon 2 of the *Soat1* gene, encoding acyl-CoA:cholesterol acyl transferase 1, also known as ACAT1, as the strongest locus modifying cholesterol ester loading[16].

The DBA/2J and AKR/J inbred strains have been useful in many areas of mouse physiology and disease. For example, the DBA/2J strain is susceptible to epicardiac calcification, which was mapped to the *Dyscalc1* QTL; and, the causal gene was identified *Abcc6* gene which has undetectable expression in the DBA/2J strain[19]. Another feature of the DBA/2J strain is that ~70% of these mice develop glaucoma by 12 months of age, after iris pigment dispersion (ipd) and iris stromal atrophy (isa)[15]. In a DBA/2JxC57BL/6J strain intercross, the ipd and isa phenotypes segregated to the *ipd* locus on chromosome 6 and the *isa* locus on chromosome 4[20]. Subsequent DBA/2JxCAST/Ei intercrosses fine mapped the *ipd* locus, and led to the discovery that this phenotype was due to a nonsense mutation in the *Gpnmb* gene (*Gpnmb^R150X^*) in the DBA/2J strain[6]. Furthermore, DBA/2 substrains with the wildtype *Gpnmb* gene do not have the ipd phenotype[6]. Thus, the same DBA/2J *Gpnmb* nonsense allele responsible for decreased macrophage lysosome function in our study is responsible for the ipd phenotype in the eye. *Gpnmb* encodes for Glycoprotein Non-Metastatic Protein B (GPNMB) which was originally discovered in a melanoma cell line[21]. This protein, also called osteoactivin, DC-HIL, or hematopoietic growth factor inducible neurokinin-1, has been studied extensively in many contexts including cancer, kidney injury, obesity, non-alcoholic steatohepatitis, Parkinson disease, osteoarthritis, lysosome storage disorders, and heart failure; and, in most of these contexts expression of GPNMB is induced by the related pathology, likely in response to lysosomal stress[12, 22–27]. However, loss of *Gpnmb* expression in DBA/2J mice is associated with preserved cardiac function after myocardial infarction[28]. Thus, there is much interest in *Gpnmb* and the role it plays in a multitude of diseases and in normal physiology.

In the current study, we verified that the *Gpnmb* null allele in the DBA/2J strain was responsible for the *Mlfm1* QTL on chromosome 6, the strongest locus for macrophage lysosome function. In addition to the *Mlfm1* QTL, we identified 3 other *Mlfm* loci on the distal region of chromosome 2 (*Mlfm3*) and on the proximal and distal regions of chromosome 17 (*Mlfm2* and *Mlfm4*). Further study will be needed to elucidate these genes and determine if any of these can account for the decreased autolysosome formation and lipid droplet turnover observed in DBA/2J macrophages.

We were able to identify the causal gene for the *Mlfm1* QTL without laborious breeding of congenic strains for fine mapping. This was aided by several factors including the use of an F_4_ intercross cohort leading to more recombinations per chromosome and by performing a high density genome scan leading to precision QTL mapping. Another fortuitous factor was the availability of the DBA/2 substrain expressing wildtype *Gpnmb*. In our experience several phenotype assays performed in mice have high coefficients of variation. For example, fatty streak aortic root lesion areas in 16 week old chow diet-fed apoE-deficient mice on inbred background strains often yield coefficients of variation approaching 50%. This large phenotypic variation, due to either stochastic or subtle environmental differences, leads to less power to detect QTLs. In the current study, we used ex-vivo cell based assays, which had smaller coefficients of variation of ~10%, leading to better power for QTL analysis even with a smaller sample size compared to many mouse based QTL studies. We call this ex vivo cell based approach ‘QTL in a dish’. One advantage of this method is the ability to treat cells with compounds or conditions that would be difficult to perform or painful in live mice. The related method ‘GWAS in a dish’ is being used to study phenotypes in different tissues derived from the differentiation of human induced pluripotent stem cells originating from cohorts consisting of ~ 100 to 200 subjects[29–31].

## Materials and Methods

### Mouse strains

AKR/J, DBA/2J, and DBA/2J-Gpnmb^+^/SjJ (stock # 007048) mice were obtained from JAX. The DBA/2J-Gpnmb^+^/SjJ mice were from the DBA/2 *Sandy* substrain, which was separated from the main DBA/2J line in the early 1980s, before the *Gpnmb^R150X^* null allele was fixed in the DBA/2J stock. Modern backcrossing to DBA/2J mice was performed to maintain the wildtype *Gpnmb* (g^+^) allele on the DBA/2J background[15]. All mouse studies were approved by the Cleveland Clinic Animal Care and Use Committee.

### Generation and genotyping of AKR/JxDBA/2J F_4_ mice

Parental male AKR/J and female DBA/2J mice were crossed to generate the F_1_ generation, fixing the Y chromosome from the AKR/J strain. Two breeding pairs of F_1_ mice were bred to generate the F_2_ mice, and two breeding pairs of F_2_ mice were used to generate F_3_ mice. Six breeding pairs of F_3_ mice were used to generate the 122 F_4_ mice, which consisted of 70 males and 52 females[16]. Healthy F_4_ mice were sacrificed at 8-10 weeks of age. Ear tissue was collected from each mouse and digested overnight at 55°C in lysis buffer containing 20 mg/mL proteinase K. DNA was ethanol precipitated and resuspended in 10 mM Tris 1 mM EDTA (pH=8). Femurs were promptly flushed after sacrifice, and bone marrow cells were washed, aliquoted, and cryopreserved. Cells were thawed and differentiated into macrophages at the time of experimentation, as described below. F_4_ mice were genotyped as described previously[16]. Briefly, the GeneSeek MegaMUGA SNP array was used, and filtering for call frequency and strain polymorphism using parental and F_1_ DNA yielded 16,975 informative SNPs that were used for QTL analysis. All marker locations are based on NCBI Mouse Genome Build 37.

### Bone marrow macrophages

Bone marrow derived macrophages were obtained from F_4_ mice and female mice on the AKR/J, DBA/2J and DBA/2g^+^ background. Bone marrow cells were suspended in macrophage growth medium (DMEM, 10% FBS, 20% L-cells conditioned media as a source of MSCF) as previously described[32, 33] and plated in tissue culture coated 6, 12, or 24 well plates. The media was renewed twice per week. Cells were used for experiments 10 to 14 days after plating when the bone marrow cells were confluent and fully differentiated into macrophages. When required, AKR/J cells were transfected with 50 nM silencer-select *Gpnmb* (4390771, Thermofisher Scientific) or control (4390843, Thermofisher Scientific) siRNA using TransIT-TKO (MIR2150, Mirus) as described by the manufacturer. Cells were incubated with the siRNA complexes for 48h, media was then replaced with fresh macrophage growth media (in the presence or absence of 50 μg/ml AcLDL as indicated), and incubated for another 24h before experiments.

### Lipoprotein preparations

Human LDL (1.019 < d < 1.063 g/mL) were prepared by ultracentrifugation from de-identified expired blood bank human plasma (exempt from human research rules as determined by the Cleveland Clinic Institutional Review Board). LDL was acetylated as described previously[34, 35] and dialyzed against PBS with 100 μM EDTA and 20 μM BHT. Protein concentrations of lipoproteins were determined using an alkaline Lowry assay[36]. When indicated, cells were loaded with 50 μg/mL of AcLDL for 24h.

### Lysosome assays

In order to determine lysosome volume, live cells were first stained with LIVE/DEAD Fixable Blue Dead Cell Stain (L23105, Thermofisher Scientific), to gate on live cells. The cells were fixed in 4% paraformaldehyde and permeabilized with saponin and lysosomes were labeled using 10 μg/mL dilution of FITC-labeled antibody against mouse Lamp-1 (ab24871, abcam), a lysosomal structural protein. To validate the use of DQ-ovalbumin as a surrogate measure of lysosome function, cells were pre-treated for 3h in absence or presence of 10 μg/mL E64d (E8640, Sigma-Aldrich) and 10 μg/mL pepstatin A (P5318, Sigma-Aldrich) before incubating for 30 min with the reagent. To measure lysosome function, macrophages were labeled with alexa647 labeled DQ-ovalbumin (A-DQ-ova). This was prepared using 1 mg of DQ-ovalbumin (D12053, Thermofisher Scientific) in 0.1 M sodium bicarbonate that was incubated with 98 μg of Alexa Fluor 647 succinimidyl ester (A20006, Thermofisher Scientific) for 1h at room temperature (3:1 dye:protein mole ratio). The reaction was stopped by incubating the conjugate with 0.1 mL of 1.5 M hydroxylamine (pH 8.5) for 1h at room temperature. The conjugate was purified by extensive dialysis. Macrophages were incubated with 2 μg/mL of A-DQ-ova for 1h, washed with PBS and suspended using CellStripper (25056CI, Corning). To evaluate lysosomal pH, cells were incubated for 18h with 1 mg/mL FITC-TAMRA dextran (D1951, Thermofisher Scientific) followed by a 4h chase period in absence or presence of 10 μM Bafilomycin A1 (B1793, Sigma-Aldrich) for the indicated times. In all experiments, 10,000 cells were analyzed by flow cytometry with a LSRII device (BD) using the following lasers and filters: 488nm excitation and 515/20nm emission (FITC and Bodipy), 639nm excitation and 660/20nm emission (Alexa647) and 532nm excitation and 575/26nm emission (TAMRA). Flowjo software was used to export data for each cell for ratiometric analyses.

### Western blot

AKR/J, AKRg^-^, DBA/2J and DBA/2g^+^ macrophages were lyzed in RIPA buffer and equal protein levels loaded on 4-20% tris-glycine gels. After transfer, membranes were probed with antibodies against GPNMB (AF2330, R&D Systems) and GAPDH (FL-335, Santa Cruz).

### Quantitative Trait Loci (QTL) analysis

QTL mapping of macrophage lysosome function (*Mlfm*) from 120 out of 122 AKR/JxDBA/2J F_4_ BMDMs (the other 2 lines did not yield viable cells) was performed using R/qtl software, with the final genotype and phenotype data formatted for analysis in the Data Supplement S1 Table[37]. The “scanone” function was utilized using Haley-Knott regression by specifying the “method” argument as “hk”. Genome-wide p-values were ascertained via permutation analysis, using 10,000 permutations by specifying the “n.perm” argument in the “scanone” function. QTL 90% confidence intervals were calculated using the 1-LOD drop off method. The credible interval for the *Mlfm1* locus was also determined by using the Bayesian credible interval (“bayesint”) function in R/qtl, with the “prob” argument set at 0.95. Since *Mlfm1* had a significantly higher peak LOD score than any other locus, QTL mapping was performed using the genotype from the best associated *Mlfm1* marker as an additive covariate (“addcovar”) in the “scanone” function of R/qtl. The *Mlfm1* corrected data were subjected to 10,000 permutation analyses to determine genome-wide p-values. To aid in prioritizing candidate genes, a custom R function termed “flank_LOD” was written (http://www.github.com/BrianRitchey/qtl). This “flank_LOD” function utilizes the “find.flanking” function in R/qtl and returns the LOD score of the nearest flanking marker for a given candidate gene position based on “scanone” output data. Genes in a QTL interval were determined by custom written R functions (“QTL_gene” and “QTL_summary”) which utilized publicly available BioMart data from Mouse Genome Build 37. A custom written R function (“pubmed_count”) which utilized the rentrez package in R was used to determine the number of PubMed hits for Boolean searches of gene name and terms of interest. Custom written R functions (“sanger_AKRvDBA_missense_genes” and “missense_for_provean”) were used to determine the number of non-synonymous mutations between AKR/J and DBA/2J in QTLs, as documented by the Wellcome Trust Sanger Institute’s Query SNP webpage for NCBIm37 (https://www.sanger.ac.uk/sanger/Mouse_SnpViewer/rel-1211). Custom written VBA subroutines (“Provean_IDs” and “Navigate_to_PROVEAN”) were used to automate PROVEAN software (http://provean.jcvi.org/seq_submit.php) queries for functional effects of missense mutations in each QTL, with rentrez functions utilized to retrieve dbSNP and protein sequence data. Ultimately, custom R code was used to generate output tables. Deleterious mutations were designated as defined by PROVEAN parameters[38]. Custom code can be found at http://www.github.com/BrianRitchey/qtl.

### Other statistics

Large data sets were tested for normal distributions and passed, thus parametric statistics were used. Comparison of two conditions was performed by two-tailed student t-test, and comparison of multiple conditions was performed by ANOVA with Tukey or linear trend posttest. All data are shown as mean ± S.D. Statistics were performed using GraphPad Prism software.

## Supporting information

Supplemental Tabke 1

## Supporting information captions

S1 Table. Lysosome function phenotypes and genotypes formatted for r/QTL analysis.

## Notes

http://www.github.com/BrianRitchey/qtl

